# Capsular K-antigen Coats Outer Membrane Vesicles of *Porphyromonas gingivalis*

**DOI:** 10.64898/2026.05.05.723094

**Authors:** Hyun Young Kim, Yeon Kyeong Ko, Hatice Hasturk, Frank C. Gibson, Manda Yu, Mary Ellen Davey

## Abstract

Periodontal disease is an inflammatory disorder that arises from dysbiosis of the subgingival microbiota, with *Porphyromonas gingivalis* acting as a keystone pathogen in the shift from health to disease. *P. gingivalis* employs multiple strategies to subvert host immune defenses, and its capsular K-antigen serves as a key virulence determinant. Here, a pre-adsorbed antiserum (pAds106) was generated by removing nonspecific antibodies using cells from a K-antigen–null mutant (W83ΔPG0106), resulting in exceptional specificity for the *P. gingivalis* K1-antigen. Immunofluorescence analysis revealed that the K-antigen preferentially coats outer membrane vesicles (OMVs), rather than attaching to the bacterial cell surface. This localization was further confirmed by ELISAs of density gradient ultracentrifuge-purified OMVs, with background signal detected in OMVs derived from K-antigen-deficient strains, non-K1-strains, and other oral Bacteroidetes. K-antigen–coated OMVs exhibited higher hydrophilicity and elicited weaker inflammatory responses compared to K-antigen–deficient OMVs, consistent with previously reported properties of encapsulated strains. Importantly, the antiserum detected K-antigen-coated OMVs in subgingival plaque from periodontal patients, suggesting that K-antigen is actively produced at diseased sites. These findings revise the prevailing view that K-antigen solely encapsulates the bacterial cell body and suggest that K-antigen–coated OMVs produced by *P. gingivalis* play distinct roles in immune evasion during periodontal disease.

## Introduction

Bacterial outer membrane vesicles (OMVs) are of fundamental importance to microbe-host signaling (1-4) and microbial biofilm matrix structure and function (5, 6). *Porphyromonas gingivalis*, a Gram-negative oral anaerobe is often regarded as the keystone pathogen in causing periodontitis through dysbiosis (7, 8) and related to many systemic diseases (9). *P. gingivalis* has evolved an extensive repertoire of evasive defense mechanisms to survive in an inflamed periodontal environment created by the host, including the ability to produce A-LPS, alter its lipid A (10, 11), synthesize sphingolipids (12), and secrete a variety of proteases and protein modifying enzymes (13). The type IX secretion system and OMV biogenesis serve as an elaborate interwoven secretion mechanism that allows *P. gingivalis* to modify its surrounding (14). Like the soil bacterium *Myxococcus xanthus* (15), *P. gingivalis* is a proteolytic degrader, it does not metabolize sugars instead it relies on peptides for growth. To break down protein into peptides for uptake, *P. gingivalis* releases high levels of protease-carrying OMVs (16). *In vitro* studies have clearly shown an OMV dense matrix surrounding *P. gingivalis* cells grown as colony biofilms. The OMVs not only appear as individual vesicles, but also long interconnected chains of OMVs (16, 17). It is reasonable to expect that any protein that enters the matrix *in vivo*, e.g., immunoglobulins, complement, and albumin; would be rapidly degraded by proteases, in particular the activity of the trypsin-like gingipains, ultimately providing protection and nutrients to the subgingival community.

Encapsulation is a well-known mechanism that protects bacteria from clearance by host immune defenses (18). K-antigens are capsular polysaccharides present on the surface of Gram-negative bacteria. For instance, a repertoire of over eighty K-antigens has been described in *Escherichia coli* (19). Encapsulation and thereby a reduction in clearance can lead to persistent survival and thereby long-term interplay between the bacterium and host. Capsules not only reduce the ability of the host effectors to gain access to the bacterial cell but also mask the cell surface and modulate the host’s response to the bacterium. This model is supported by studies showing that bacteria that cloak themselves in a capsule have an advantage in immune evasion (20).

Synthesis of K-antigen is a key virulence determinant of *P. gingivalis* (21). Strains of *P. gingivalis* that produce K-antigen are more resistant to phagocytosis (22) and disseminate from the site of infection in a murine lesion model (23). In contrast, *P. gingivalis* strains that are K-antigen minus adhere more readily to cultured primary gingival epithelial cells and cause a localized abscess (24). In addition, a K-antigen null mutant strain was shown to be a more potent inducer of cytokine production in human gingival fibroblasts, indicating a role for this polysaccharide in cloaking *P. gingivalis* against innate immune responses (21). Recent studies showed that infections with strains that produce K-antigen (K1 and K2 serotypes) can induce neuroinflammation, astrogliosis, cognitive decline, and histopathological signs of Alzheimer’s disease in the hippocampus of rats (25), which aligns with earlier studies showing that purified capsular polysaccharide (in particular K1 and K2 serotypes) elicit chemokine production from phagocytic cells, suggesting that the host response to this antigen may contribute to the formation of the inflammatory cell lesion observed during *P. gingivalis*-elicited periodontal disease. Notably, a recent clinical study found that among K1–K7 serotypes, the K1 capsular antigen was significantly associated with moderate to severe depressive symptoms in pregnant women (26).

In our previous studies, K-antigen detection was based on an ELISA using polysaccharide-detection enriched IgM antiserum that was generated against W83 whole cells (27-29). As the antiserum contains various types of IgM, additional steps like autoclaving are required to remove proteins and other polysaccharides which make applications like immunofluorescent staining not feasible. Recently, we developed a method to purify IgM by pre-adsorption of the antiserum with ΔPG1881 cells. We showed that the pre-adsorbed antiserum only targets the O-glycan that anchored to OMV-targeted PG1881 (30). Deletion of PG0106, a glycosyl-transferase causes the complete removal of K-antigen biosynthesis but no major impact on other surface antigens (27, 31, 32). We, therefore, in this study, utilized W83ΔPG0106 mutant to generate the pre-adsorbed antiserum (pAds106-IgM) for K-antigen detection. Based on the approach, we found that the large capsular K-antigen polysaccharide in the W83 strain is preferentially exported to OMVs. In addition, we determined that *P. gingivalis* K-antigen–coated OMVs are detected in subgingival plaque from diseased sites. These findings revise the traditional view that K-antigen functions solely as a capsule surrounding the bacterial cell. Given their small size, K-antigen–coated OMVs may disseminate to distant sites in the body and play previously unrecognized roles in systemic disease pathogenesis.

## Results

### Pre-adsorbed antiserum demonstrates specific detection of the K-antigen

IgM-enriched antiserum for capsular polysaccharide (CPS) detection was generated by multiple injections and a shortened immunization schedule (Fig. 1A) (27) and was previously shown to detect and quantify K-antigen CPS (27-29). The immunoblot using the non-pre-adsorbed antiserum showed a dominant signal above 250 kDa (Fig. 1B), a distinct feature of CPS (33) separated apart from other smaller polysaccharides associated with A-LPS or O-LPS (34). While the K-antigen signal is dominant, IgM recognizing non–K-antigen epitopes at lower molecular weights may confound interpretation if the antiserum is applied directly. Thus, autoclaving the substrate is necessary to remove heat-sensitive antigens, limiting the use of this antiserum in techniques such as immunoblotting and immunofluorescent staining. To address this limitation, we employed our recently developed pre-adsorption approach (30) to remove IgMs that target non–K-antigen epitopes (Fig. 1A). This purification strategy was implemented by pre-adsorbing the antiserum with the ΔPG0106 mutant, which has similar protein and LPS profiles (Fig. 1C & S1A) but lacks the core K-antigen biosynthesis gene, PG0106 (31). The pre-adsorbed antiserum (pAds106) was the used as the primary antibody, and anti-rabbit-IgM–HRP antibodies were used as the secondary antibodies (i.e. pAds106-IgM) in immunoblot analyses of the parent strain W83, the ΔPG0106 mutant, and two additional *P. gingivalis* strains (ATCC33277 and 381) that do not produce the K-antigen. A high–molecular-weight signal (>250 kDa) was observed exclusively in W83, with no detectable signals in the other three strains and a lower background than the non-pre-adsorbed antiserum (Fig 1C). CPSs from W83 and W83ΔPG0106 strains were then extracted using a phenol–water–based method (33) for immunoblot analysis. The CPS extracts were further treated with proteinase K to remove residual proteins and with acetic acid to remove LPS molecules (35). The high– molecular-weight signals remained strongly detectable, confirming that pAds106-IgM specifically recognizes the K-antigen CPS (Fig S1C & S1D).

**Figure 1.**
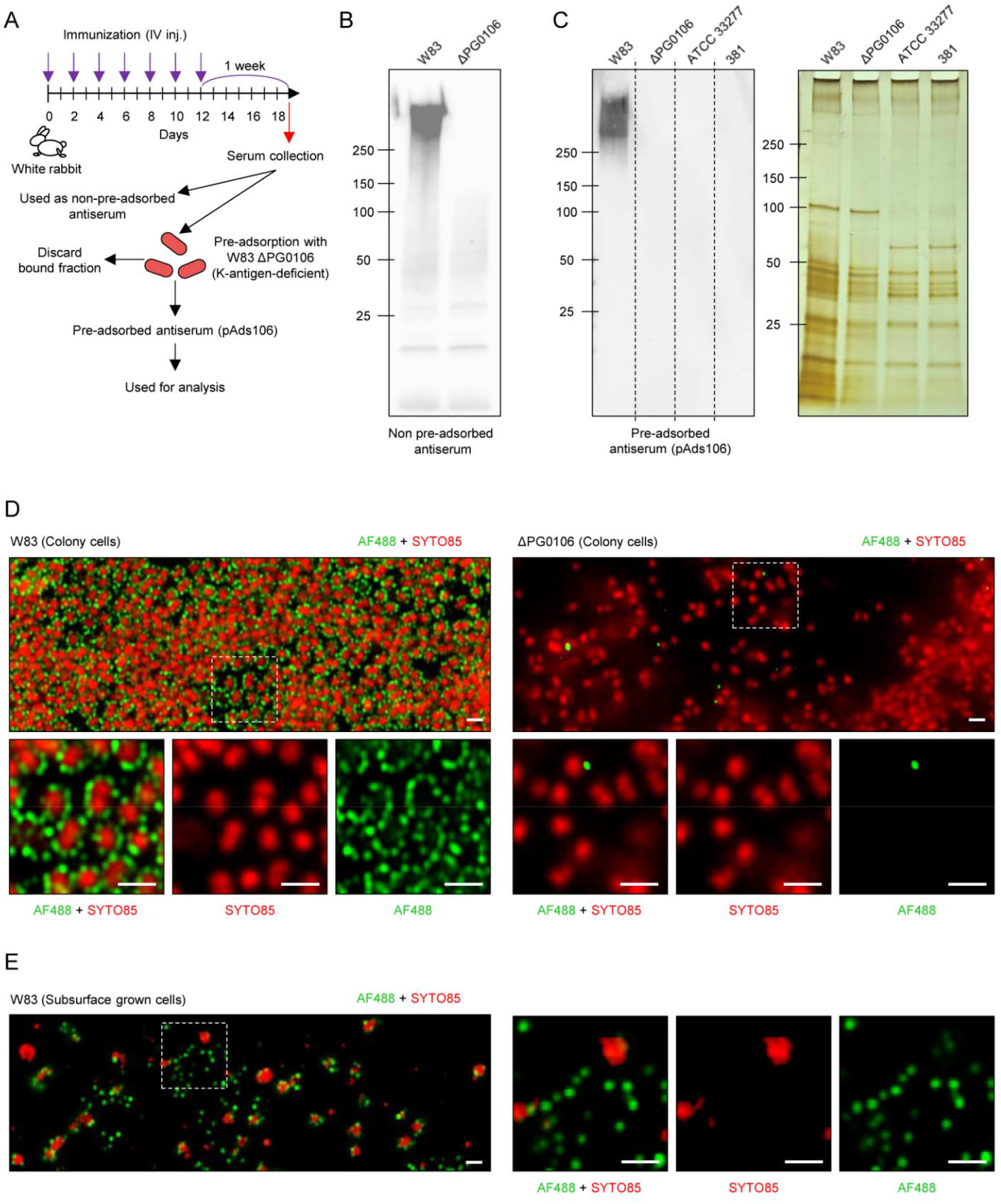
**(A)** Schematic illustration for the generation of pAds106 antiserum. **(B)** Immunoblot of cell lysates from *P. gingivalis* WT W83 and Δ PG0106 using non-pre-adsorbed antiserum as primary antibodies and anti IgM-HRP as the secondary antibodies. **(C)** Immunoblot of cell lysates from *P. gingivalis* WT W83, W83ΔPG0106, ATCC33277 and 381 strains using pAds106 antiserum as primary antibodies and anti IgM-HRP as the secondary antibodies. The samples were stained with silver stain for loading comparison. **(D)** Immunofluorescence staining WT W83 and W83ΔPG0106 colony cells using pAds106 antiserum as the primary antibodies and anti-IgM-AF488 (green) as the secondary antibodies; counterstain with and SYTO85 (red). Scale bar, 1 μm. **(E)** Immunofluorescence staining of WT W83 subsurface-grown cells using pAds106 antiserum as the primary antibodies and anti-IgM-AF488 (green) as the secondary antibodies; counterstain with and SYTO85 (red). Scale bar, 1 μm.

### Capsular K-antigen is exported to the outer membrane vesicles

As the subcellular localization of K-antigen in *P. gingivalis* has not been previously resolved using specific labeling, immunofluorescence staining of colony cells was first performed using pAds106 antiserum as the primary antibodies and Alexa Fluor 488 (AF488) conjugated anti-IgM secondary antibodies. Strong fluorescence signals were detected surrounding the WT bacterial cells but not on the ΔPG0106 mutant cells (Fig. 1D), indicating the specificity of the detection of K-antigen. Unexpectedly, rather than forming a continuous layer around the bacterial cells, the fluorescence signals appeared as discrete spherical puncta (Fig. 1D), resembling OMVs in size and abundance. Since OMVs are likely to be retained on the cell surface of *P. gingivalis* cells, to enhance visibility, a subsurface translocation growth condition (36) was employed to promote OMV release from the cell surface. Immunofluorescence assays of subsurface-grown cells revealed that the signals were predominantly associated with OMVs (Fig. 1E & S2). To further validate the microscopic observations, OMVs from W83 and the ΔPG0106 mutant strain were purified by density gradient ultracentrifugation, and fraction #5 was collected (Fig. 2A), and their physical properties were analyzed by nanoparticle tracking analysis (NTA) (Fig. 2B-2D). Across three independent OMV preparations (Fig. S3), ΔPG0106 OMVs were smaller than those from the parent strain (Fig. 2B and 2C), while no differences were observed in particle-to-protein ratio (Fig. 2D). CPSs are hydrophilic and alter the physical properties of encapsulated objects that can be assessed by bacterial adherence to hydrocarbons (BATH) assay (37, 38). Hydrocarbons such as xylene preferentially retain the hydrophobic (i.e. non-encapsulated) objects in the organic layer, resulting in fewer objects remaining in the aqueous phase. OMVs derived from W83 consistently showed lower hydrophobicity than those from the ΔPG0106 mutant, indicating the hydrophilic CPS is likely present on W83 OMVs (Fig. 2E & S3). Alcian blue staining (39) further confirmed the physical differences between W83 and ΔPG0106 OMVs, revealing a high-molecular-weight CPS signal solely in OMV extracts from the parent strain W83 (Fig. 2F). ELISA preserves antigen structure is considered more suitable for detecting native epitopes, while also allowing quantitative assessment of non-specific binding compared to immunoblot analyses. Thus, ELISAs based on pAds106-IgM were performed to analyze the specific detection of K-antigen on the OMVs. Planktonic-grown cells were washed twice by centrifugation to remove unattached OMVs. As shown in Fig. 2G and 2H, both particle- and protein-normalized consistently showed significantly higher pAds106-IgM binding in purified OMVs than in the washed planktonic-grown cells, indicating that K-antigen is preferentially associated with OMVs. Additional ELISAs were performed using OMVs purified from *P. gingivalis* strains W83, ΔPG0106 mutant 381 and ATCC33277; and three oral Bacteroidetes bacteria, *Porphyromonas endodontalis, Tannerella forsythia* and *Prevotella intermedia* (Fig. 2I and 2J). The OMVs from ΔPG0106 mutant, *P. endodontalis, T. forsythia* and *P. intermedia* show only background levels of binding to pAds106-IgM. The antiserum did not recognize CPS from OMVs or cells of non–K1-antigen serotype strains (K3: A7A1-28; K4: ATCC49417) (40) (Fig. 2L and 2M, S1B) that contain the core PG0106 gene (41). Interestingly, non-K1 *P. gingivalis* strains (Fig. 2I and 2K) exhibited low-level detection by pAds106-IgM, indicating that these strains may synthesize an alternative OMV-associated polysaccharide that requires the glycosyl-transferase PG0106. Together, these results demonstrate that capsular K1-antigen from strain W83 is preferentially exported to the OMVs in *P. gingivalis*, where it contributes to distinct physicochemical properties and can be specifically detected by pAds106-IgM.

**Figure 2.**
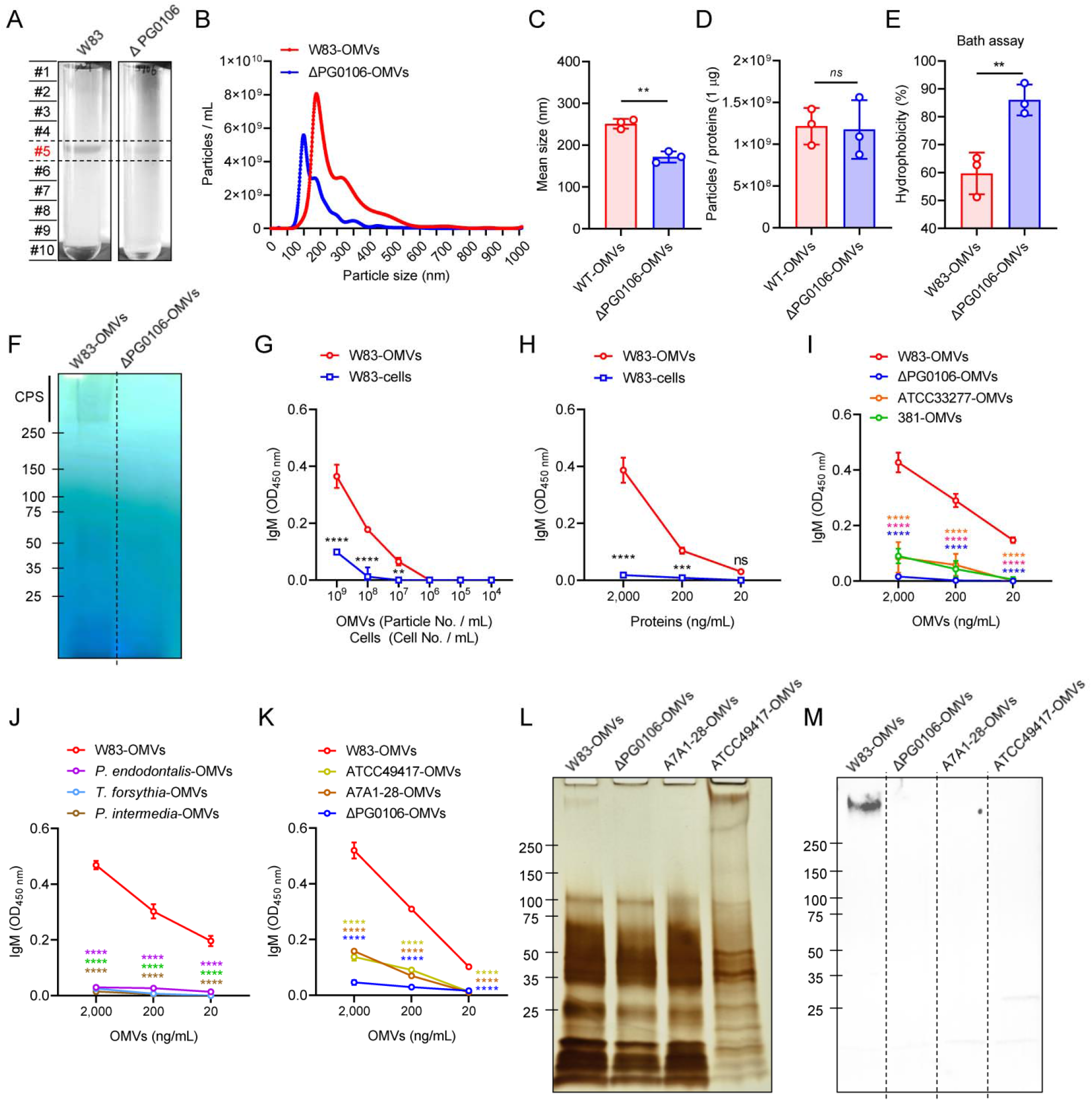
**(A)** Representative image showing the purification of *P. gingivalis* OMVs by buoyant density gradient ultracentrifugation using OptiPrep reagent. **(B, C)** Size distributions and mean particle size of W83-OMVs and ΔPG0106-OMVs were determined by NTA. **(D)** Particle-to-protein ratios (particles per 1 μg protein) were determined by NTA for particle quantification and by BCA assay for protein measurement. **(E)** Surface hydrophobicity of OMVs measured by the BATH assay. Percentage of OMVs retained after xylene treatment were quantified by NTA. **(F)** OMV lysates (50 μg of protein content) of W83 and ΔPG0106 were separated by SDS-PAGE, followed by Alcian blue staining. **(G)** ELISA using pAds106-IgM, in which W83 cells and OMVs were quantified based on particle number, and **(H)** ELISA in which W83 cells and OMVs were normalized by protein concentration. **(I)** ELISA using pAds106-IgM to detect OMVs from W83, ΔPG0106, ATCC33277, and 381 strains, **(J)** other oral Bacteroidetes species-OMVs (*P. endodontalis, T. forsythia*, and *P. intermedia*), and **(K)** *P. gingivalis* strains with different K serotypes (A7A1-28 and ATCC49417). Three independent OMV preparations were analyzed. Statistical significance was determined by one-way ANOVA with Bonferroni’s multiple comparison test (**C, D, E**) or two-way ANOVA with Bonferroni’s multiple comparison test (**G, H, I, J, K**). Statistical significance was indicated by lines connecting the compared groups; for ELISA, each sample was compared only to W83-OMVs at matched concentration. \**P* < 0.05, \*\**P* < 0.01, \*\*\**P* < 0.001, \*\*\*\**P* < 0.0001. *ns* denotes not significant. OMV lysates (10 μg of protein content) of W83, ΔPG0106, ATCC49417, and A7A1-28 were separated by SDS-PAGE, followed by silver staining **(L)** and immunoblotting using pAds106-IgM **(M)**.

### Detection of K-antigen in the periodontitis dental plaque

Following the biochemical and *in vitro* findings, pAds106-IgM was used to detect K-antigen polysaccharides in subgingival plaque specimens. To first confirm the presence of *P. gingivalis* in these samples, immunofluorescence staining was performed using Alexa Fluor 647 conjugated anti-*P. gingivalis* antibodies (Anti-Pg-AF647; red), with SYTO9 (green) as a nucleic acid counterstain (Fig. 3A). *P. gingivalis* was detected, with antibody signals localized around the cell periphery. Next, pAds106 was applied as the primary antibody, and Alexa Fluor 488–conjugated anti-IgM (pAds106-IgM-AF488; green) was used as the secondary antibody, with SYTO85 (red) as a nucleic acid counterstain. In contrast to the anti-*P. gingivalis* antibodies, punctate fluorescence signals were observed attached to the bacterial cells as well as detached from the cells (Fig. 3B & S4). To confirm that these punctate signals were associated with *P. gingivalis*, double staining was performed using anti-Pg–AF647 (red) and pAds106-IgM–AF488 (green)(Fig. 3C). The results showed co-localization of the punctate signals with *P. gingivalis* cells that were similar with the *in vitro* results (Fig. 1D). Notably, not all *P. gingivalis* cells were surrounded by K-antigen signals, suggesting either the presence of non-K1 strains in the sample or that most OMVs are released from the bacterial surface. In contrast, a plaque sample from a healthy individual showed reduced *P. gingivalis* abundance and minimal K-antigen–associated OMV signals (Fig. S5). These results indicate that K-antigen–carrying OMVs are present in periodontal plaque samples *in situ*.

**Figure 3.**
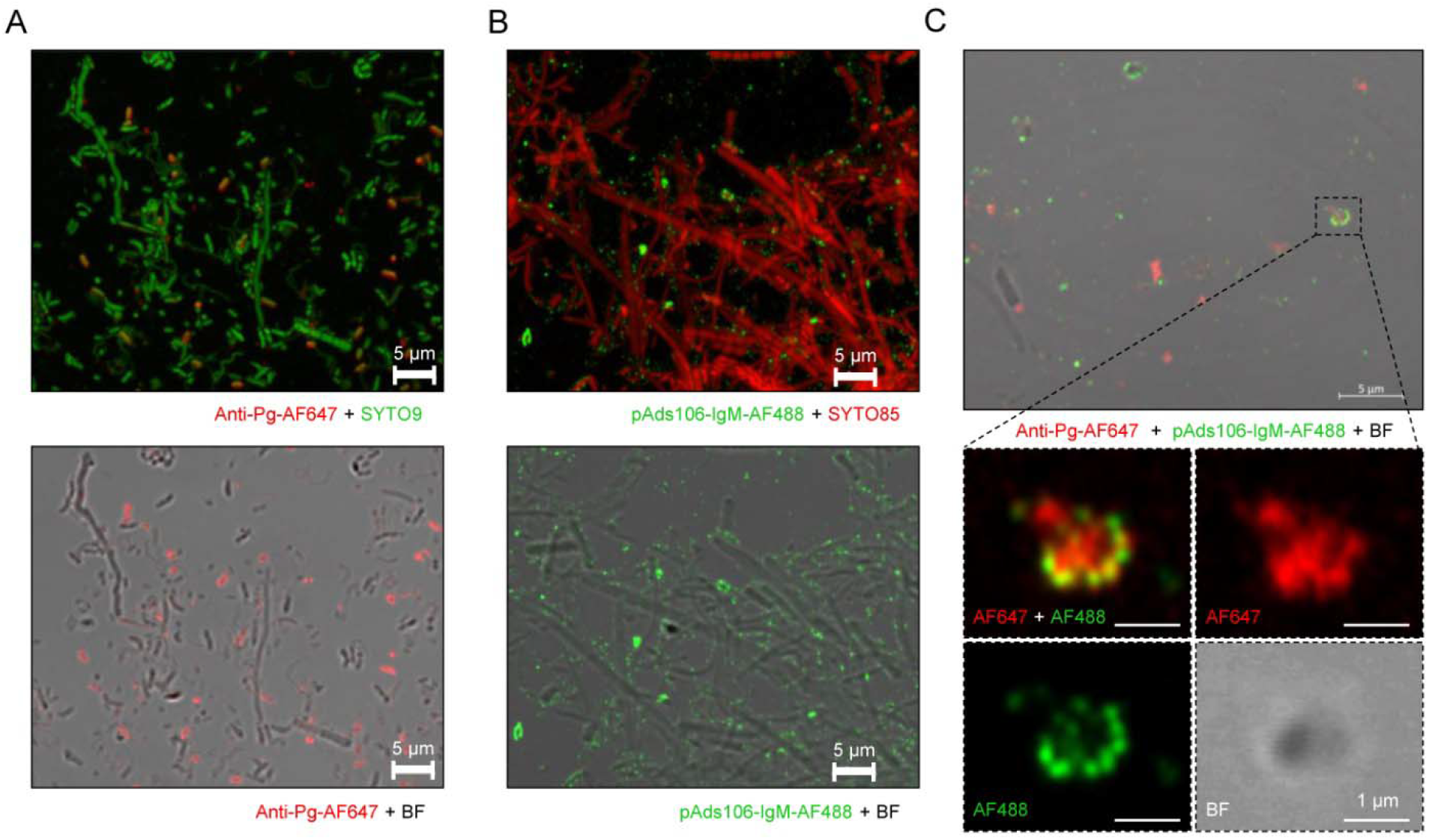
**(A)** Immunofluorescence staining of periodontal plaque sample using AF647 conjugated anti-*P. gingivalis* antibodies (Anti-Pg-AF647; red); counterstain with and SYTO9 (green). **(B)** Immunofluorescence staining of periodontal plaque sample using pAds106 as the primary antibodies and anti-IgM-AF488 as the secondary antibodies (green); counterstain with and SYTO85 (green). **(C)** Periodontal plaque samples were double-stained by sequential incubation with pAds106 primary antibody and anti-IgM–AF488 secondary antibody (green), followed by staining with anti-Pg–AF647 antibodies (red).

### K-antigen– deficient OMVs differentially regulate host inflammatory responses

Non-encapsulated *P. gingivalis* strains have been shown to elicit higher inflammatory responses than their encapsulated counterparts in multiple host cell types, including human gingival fibroblasts and murine innate immune cells, while capsulation contributes to bacterial survival and virulence (22). To determine whether this difference is mediated by encapsulated OMVs, a Transwell co-culture system was employed. THP-1 human macrophage-like cells were seeded in the lower chamber, while *P. gingivalis* W83 or ΔPG0106 cells were added to the upper chamber, which permits the passage of small particles such as OMVs but prevents direct bacterial contact. The results showed that both strains induced the production of TNF-α, IL-1β, and IL-8; however, the ΔPG0106 mutant elicited significantly higher cytokine levels than the parent strain (Fig. 4A). When purified OMVs were directly applied to THP-1 cells, a similar trend was observed, with ΔPG0106 OMVs inducing higher cytokine production than W83 OMVs (Fig. 4B). A similar experimental setup was performed using telomerase-immortalized gingival keratinocytes (TIGK cells), a model for oral gingival epithelium. In the Transwell system, both W83 and the corresponding ΔPG0106 mutant suppressed IL-8 production, with W83 exhibiting significantly stronger suppression (Fig. 4C). Consistently, W83-derived OMVs also showed a higher inhibitory effect on IL-8 production compared to ΔPG0106 OMVs (Fig. 4D). These findings indicate that encapsulated OMVs recapitulate the immunomodulatory properties previously observed in whole-cell studies. To exclude the possibility that these differences were due to variation in gingipain activity between W83 and ΔPG0106 OMVs, recombinant TNF-α, IL-1β, and IL-8 were incubated with OMVs from both strains. No significant differences in cytokine degradation were observed (Fig. 4E), suggesting that the differential immune responses are not attributable to gingipain-mediated proteolysis.

**Figure 4.**
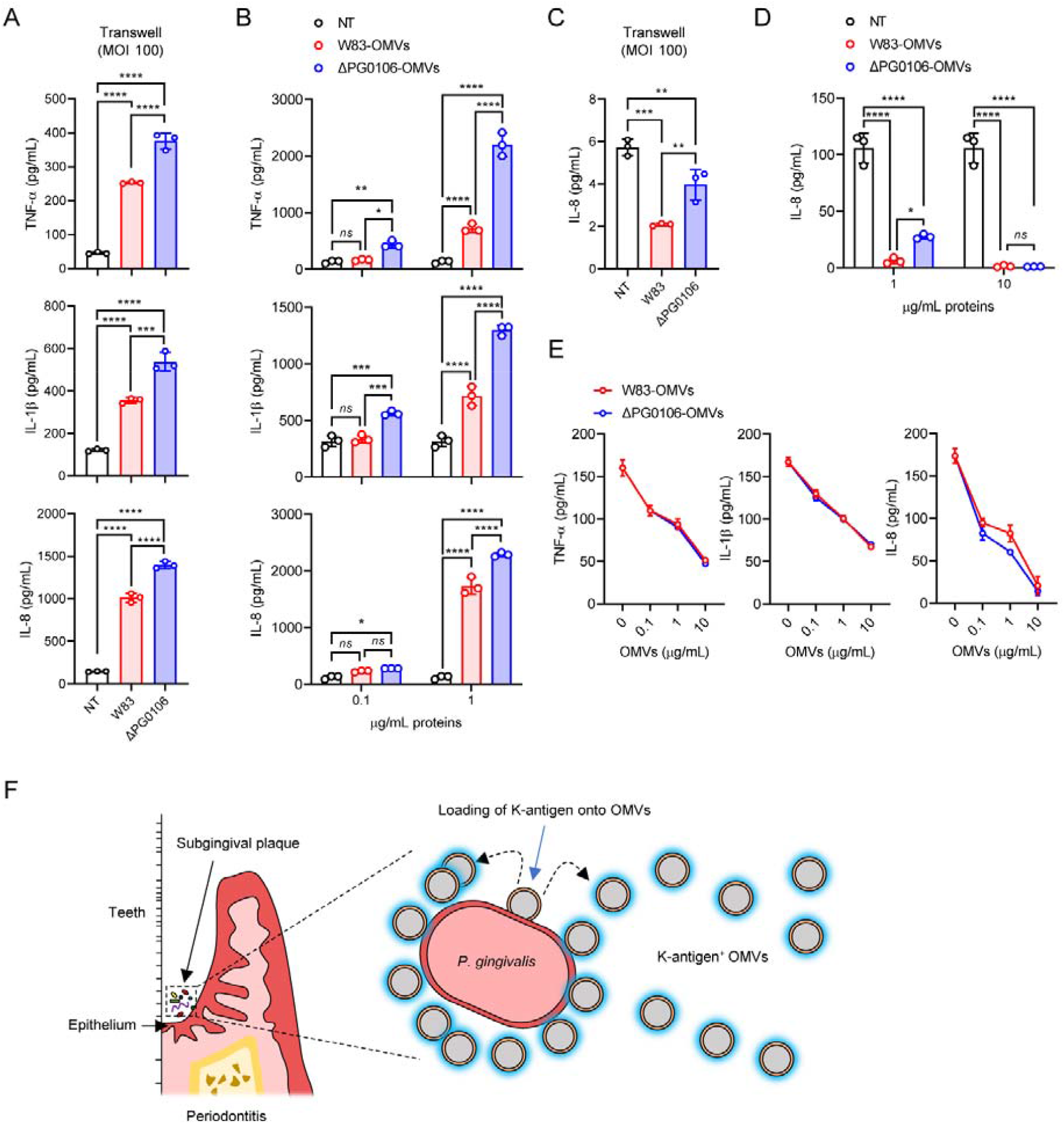
**(A)** THP-1 cells (1 × 10^6^ cells/well) were seeded in the lower chamber of a Transwell plate, and *P. gingivalis* W83 or W83ΔPG0106 (MOI 100) were added to the upper chamber in antibiotic-free medium, followed by incubation for 24 h. **(B)** THP-1 cells (1 × 10^6^ cells/well) were treated with W83-OMVs or ΔPG0106-OMVs at 0.1 or 1 μg/mL for 24 h. **(C)** TIGK cells (1 × 10^5^ cells/well) were seeded in the lower chamber of a Trans-well plate, and W83 or ΔPG0106 (MOI 100) were added to the upper chamber in antibiotic-free medium. **(D)** TIGK cells (1 × 10^5^ cells/well) were treated with W83-OMVs or ΔPG0106-OMVs at 1 or 10 μg/mL for 24 h. **(E)** Recombinant TNF-α, IL-1β, and IL-8 (200 μg/mL) were incubated with W83-OMVs or ΔPG0106-OMVs at the indicated concentrations (0, 0.1, 1, and 10 μg/mL) for 24 h at 37°C. Following incubation, supernatants were collected and used to measure cytokine levels by ELISA. Statistical significance was determined by one-way ANOVA with Bonferroni’s multiple comparison test **(A, C)** or two-way ANOVA with Bonferroni’s multiple comparison test **(B, D)**. Statistical significance was indicated by lines connecting the compared groups, with \**P* < 0.05, \*\**P* < 0.01, \*\*\**P* < 0.001, \*\*\*\**P* < 0.0001. ns denotes not significant. **(F)** Schematic illustration of the detection of K-antigen–positive *P. gingivalis* OMVs in subgingival plaque samples.

## Discussion

The major finding of this study is that K-antigen polysaccharide produced by *P. gingivalis* W83, which is hydrophilic and heat stabile is exported and attached to the OMVs rather than encapsulates the bacterial cell body, as has been widely acknowledged in many bacterial species. This unexpected observation prompted us to employ multiple approaches to verify this finding. The data show that detection of K-antigen with pAds106-IgM is robust and specific to CPS. Both immunoblots and ELISAs determined that pAds106-IgM has little if any non-specific binding to strains that lack K-antigen (381, ATCC33277 and the W83ΔPG0106 mutant). We also ruled out the binding of pAds106-IgM to proteins and the LPS of this organism. Remarkably, K-antigen-deficient OMVs demonstrate variations in the physical properties and immunological related properties. When compared to the K-antigen null OMVs, W83 OMVs are more hydrophilic and elicit a weaker immune response, matching data of previous studies using whole *P. gingivalis* cells (40). The earlier understanding that CPS encapsulated the cell bodies is attributed to the limited specificity and resolution of earlier observations, which relied largely on the nonspecific detection of a colorless halo surrounding the cells by India ink staining (31, 32) or on the apparent thickness of the signals surrounding the cells observed by electron microscopy (27). Because *P. gingivalis* produces a large quantity of OMVs that envelop the cell body (16, 17), the K-antigen present on these OMVs is likely to function similarly to the cell-surface capsule, repelling India ink and generating the colorless halo that has been observed. In contrast, high resolution Airyscan microcopy (42) combined with specific pAds106-IgM used in this study was able to clearly observe and delimit the OMV structures on the cell surface.

K-antigen producing encapsulated strains are considered to be more virulent, i.e., able to subvert inflammatory responses (22). Consistent with previous findings with *P. gingivalis* cells, K-antigen–containing OMVs elicited weaker inflammatory responses in human macrophages and oral keratinocytes compared to K-antigen–deficient OMVs, supporting a conserved immune-modulatory role of K-antigen in dampening host inflammation. Given that *P. gingivalis* OMVs carry multiple Toll-like receptor (TLR) ligands and virulence factors capable of activating host innate immune signaling (43), our findings suggest that K-antigen on OMVs functions similarly to capsular structures on bacterial cells by masking underlying immunostimulatory components and thereby attenuating host responses. Intriguingly, pAds106-IgM exhibited high specificity toward the K1 antigen subtype, with no detectable reactivity against K3 or K4 antigens tested in this study. Notably, the K1 subtype has been more strongly associated with virulence across *in vitro, in vivo*, and clinical studies (26, 40). This finding echoes the immunofluorescence staining of subgingival plaque samples, in which K1 antigen was detected on OMVs, raising the possibility that K-antigen–bearing OMVs, particularly those expressing K1, may disseminate systemically via the bloodstream and contribute to disease pathogenesis at distal sites. We therefore envision that pAds106-IgM could also serve as a useful tool for the detection and neutralization of *P. gingivalis* OMVs, thereby helping to mitigate associated systemic diseases. Furthermore, it will be fascinating to elucidate the mechanism behind K-antigen CPS export onto OMVs and to determine whether similar capsular antigen coating occurs in OMVs from other bacterial species.

## Materials and Methods

### Bacterial strains and culture conditions

*P. gingivalis* strains were grown on Trypticase Soy Agar plates supplemented with 5 μg /ml hemin, 1 μg/ml menadione, and 5 % defibrinated sheep blood (Northeast Laboratory Services) (BAPHK) at 37°C in an anaerobic chamber (Coy Lab Products) with an atmosphere containing 5 % hydrogen, 10 % carbon dioxide, and 85 % nitrogen. Planktonic cultures of all bacterial strains were grown in Trypticase Soy Broth (Becton, Dickinson and Company) supplemented with 5 μg/ml hemin and 1 μg/ml menadione (TSBHK). In this study, *Porphyromonas. endodontalis* ATCC35406, *Prevotella intermedia* strain 17 and *Tannerella forsythia* ATCC43037 were used as the represented oral Bacteroides. Subsurface growth conditions were performed as previously described (30, 36, 44) with modifications. Detailed procedures are provided in the Supplementary Methods.

### Clinical samples

Subgingival biofilm samples were collected from volunteers who provided informed consent prior to study participation, in accordance with a study protocol approved by the ADA Forsyth Institutional Review Board (Protocol #18-06, approval date: 08May2025). Participants’ eligibility (e.g., medically healthy, non-smokers, no antibiotic use in the past 6 months, etc.) and oral and periodontal health were assessed using standard protocols on a separate day prior to sampling. At the sampling visit, participants presented to the clinic without having performed their morning oral hygiene (brushing, flossing, or mouthwash use) and without having eaten or drunk for 2 hrs. Sites were isolated with cotton rolls, and supragingival plaque was gently removed with sterile Gracey curettes. Subgingival biofilm samples were then obtained from a site with ≥6 mm pocket depth and presence of bleeding on probing in individuals with periodontal disease (Stage III, Grade B, N=2) and from a site with ≤3 mm pocket depth and no bleeding on probing in a healthy control individual. Samples were placed in individual microcentrifuge tubes (Eppendorf tubes, Thermo Fisher Scientific) containing 4% paraformaldehyde (PFA) and transferred to the laboratory for processing and assays.

### Pre-adsorption of antiserum with *P. gingivalis* ΔPG106 cells

Pre-adsorption method was described previously (30) with modifications. *P. gingivalis* ΔPG0106 cells were harvested from BAPHK plates, resuspended in Phosphate-buffered saline (PBS), and adjusted to 70% ethanol for a 30-min incubation. The cells were then centrifuged and resuspended in PBS to an OD_600_=3. In the pre-adsorption mixture, 200 μl of ethanol treated cells, 200 μl of anti-W83 antiserum were mixed in 0.8 ml of PBS with protease inhibitors (Halt protease inhibitor, Thermo Fisher Scientific) and incubated at room temperature for 30 mins. The cells were removed by centrifugation at 12,000 x *g* for 3 mins. Fresh 200 μl of ethanol treated cells was added to the supernatant and repeated the binding step. The pre-adsorbed serum was mixed with 1ml of glycerol (50% final concentration) and stored at −2°C.

### Outer membrane vesicle (OMV) purification

Bacterial strains for OMV purification were cultured at 37°C in an anaerobic chamber for 16 hrs in TSBHK as previously described (30). After culturing, the optical density (OD_600_) was adjusted to 1.0–1.2. OMVs were purified as described previously with minor modifications (45, 46). The size distribution and particle concentration of OMVs were determined by nanoparticle tracking analysis (NTA) using NanoSight Pro (Malvern Panalytical, Model no. HBG5000). Detailed procedures are provided in the Supplementary Methods.

### Human cell line experiments

THP-1 cells and human telomerase-immortalized gingival keratinocytes (TIGK) were cultured and used for Transwell assays as previously described (47). Cells were treated with live *P. gingivalis* or OMVs for 24 hrs, and TNF-α, IL-1β, IL-6, and IL-8 levels were measured using R&D DuoSet ELISA kits according to manufacturer’s instructions.

### Cytokine degradation assay

Cytokine degradation by OMVs was determined as previously described (48). Briefly, 200 μg/mL recombinant cytokines (R&D Systems, Minneapolis, MN, USA) were incubated with *P. gingivalis* OMVs at the indicated concentrations for 24 hrs at 37°C in a humidified CO_2_ incubator, and the remaining cytokines were quantified by ELISA. Cytokines incubated under the same conditions without OMVs were used as a negative control.

### Immunoblot analysis

For K-antigen detection, cells (OD_600_ = 1, in PBS) or OMVs corresponding to 10 μg of total protein were resuspended in an equal volume of 4X Laemmli sample buffer (Bio-Rad) and boiled for 5 mins. Proteinase K was added to remove protein if necessary (28). Detailed procedures are provided in the Supplementary Methods.

### Immunofluorescent staining and confocal microscopy, enzyme-linked immunosorbent assay (ELISA), capsular polysaccharide extraction, Alcian blue staining and BATH assay

Detailed procedures are provided in the Supplementary Methods. The immunofluorescent microscopy was based on our previous studies (30, 49) with modifications. A direct ELISA using anti-*P. gingivalis* serum was conducted as described previously (50) with modifications. Bacterial adherence to hydrocarbons (BATH) assay was performed based on previously described methods (37) with modifications for OMV analysis.

## Acknowledgements

We thank Dr. Batbileg Bor and Dr. Deepak Chouhan (ADA Forsyth Institute) for providing TIGK cells, and Dr. Jose Solbiati (University of Florida) for providing the *P. gingivalis* A7A1-28 and ATCC 49417 strains. This work was supported by the National Institute of Dental and Craniofacial Research of the National Institutes of Health under award numbers R01DE033659 to M.E.D and R01DE031159 awarded to M.E.D. and F.C.G.

## Notes

### Competing Interest Statement

The authors have declared no competing interest.

