## Supplemental Information for "Capsular K-antigen Coats Outer Membrane Vesicles of *Porphyromonas gingivalis*"

**Supporting Information (Supplementary Figures and Supplementary Methods)**

**Figure S1.**

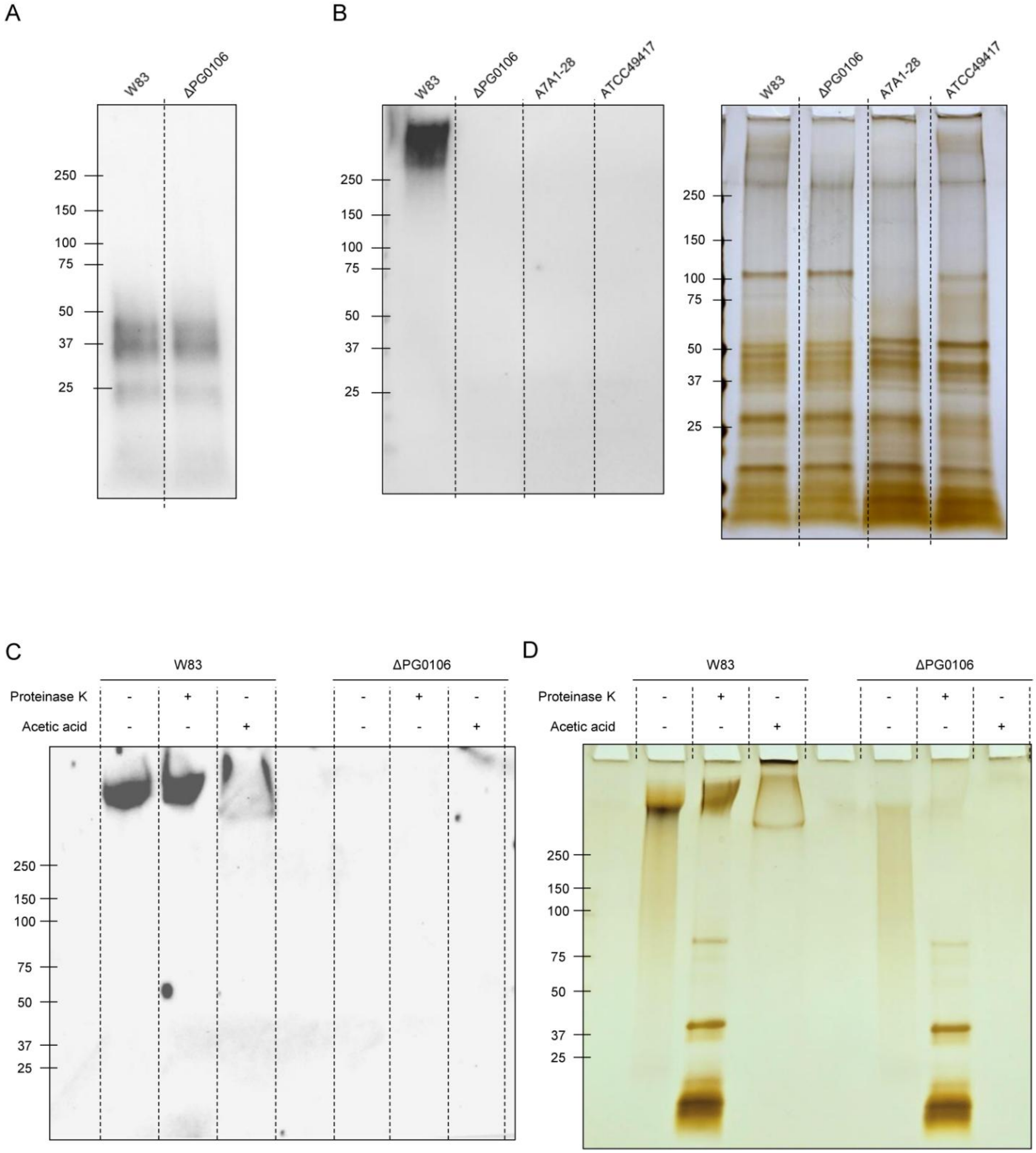


**Figure S1. (A)** Immunoblot of colony-derived cells from WT W83 and ΔPG0106 mutant probed with monoclonal IB5 antibodies (Bainbridge et al., 2015), followed by anti-mouse IgG–HRP for detection of A-LPS. **(B)** Immunoblot and silver stain of colony-derived cells from WT W83, ΔPG0106 mutant, A7A1-28 and ATCC49417. pAds106 was used as the primary antibodies, followed by anti-rabbit-IgM-HRP. **(C & D)** Immunoblot and silver stain of purified CPS from WT W83 and ΔPG0106 mutant. CPS was further treated with proteinase K and acetic acid for removal of proteins and LPS respectively. pAds106 was used as the primary antibodies, followed by anti-rabbit-IgM-HRP.

**Figure S2.**

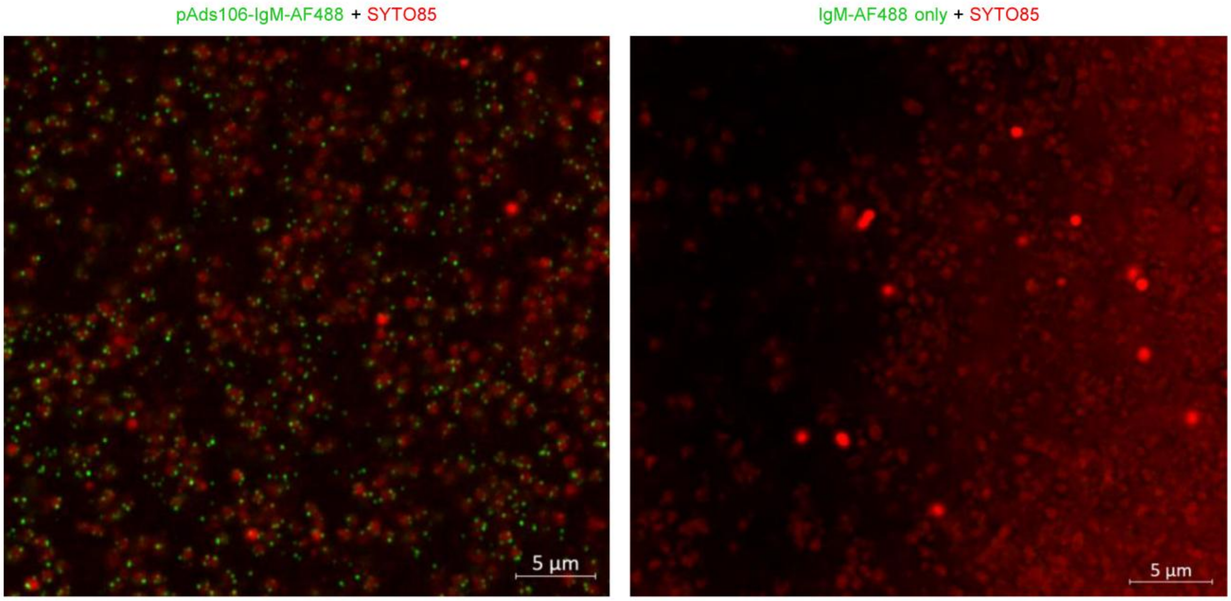


**Figure S2.** A negative control of subsurface grown cells subjected to centrifugation (right) was stained with secondary antibody IgM–AF488 alone, in comparison with the positive signal obtained using pAds106 as the primary antibody (left). Both samples were counterstained with SYTO85 (red).

**Figure S3.**

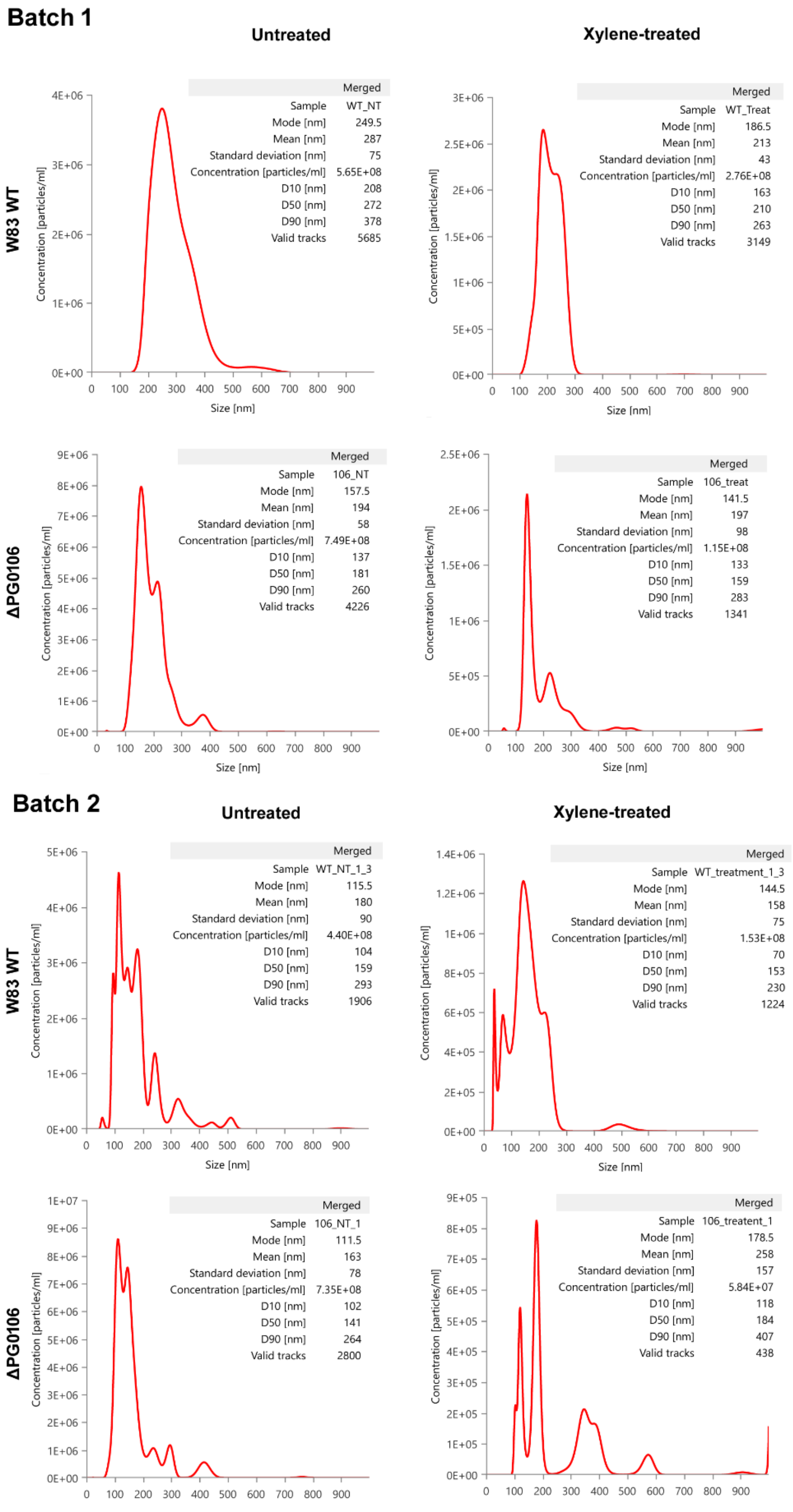
Batch 1
Untreated
Xylene-treated
W83 WT
ΔPG0106
Merged
Sample WT_NT
Mode [nm] 249.5
Mean [nm] 287
Standard deviation [nm] 75
Concentration [particles/ml] 5.65E+08
D10 [nm] 208
D50 [nm] 272
D90 [nm] 378
Valid tracks 5685
Merged
Sample WT_Treat
Mode [nm] 186.5
Mean [nm] 213
Standard deviation [nm] 43
Concentration [particles/ml] 2.76E+08
D10 [nm] 163
D50 [nm] 210
D90 [nm] 263
Valid tracks 3149
Merged
Sample 106_NT
Mode [nm] 157.5
Mean [nm] 194
Standard deviation [nm] 58
Concentration [particles/ml] 7.49E+08
D10 [nm] 137
D50 [nm] 181
D90 [nm] 260
Valid tracks 4226
Merged
Sample 106_treat
Mode [nm] 141.5
Mean [nm] 197
Standard deviation [nm] 98
Concentration [particles/ml] 1.15E+08
D10 [nm] 133
D50 [nm] 159
D90 [nm] 283
Valid tracks 1341
Batch 2
Untreated
Xylene-treated
W83 WT
ΔPG0106
Merged
Sample WT_NT_1_3
Mode [nm] 115.5
Mean [nm] 180
Standard deviation [nm] 90
Concentration [particles/ml] 4.40E+08
D10 [nm] 104
D50 [nm] 159
D90 [nm] 293
Valid tracks 1906
Merged
Sample WT_treatment_1_3
Mode [nm] 144.5
Mean [nm] 158
Standard deviation [nm] 75
Concentration [particles/ml] 1.53E+08
D10 [nm] 70
D50 [nm] 153
D90 [nm] 230
Valid tracks 1224
Merged
Sample 106_NT_1
Mode [nm] 111.5
Mean [nm] 163
Standard deviation [nm] 78
Concentration [particles/ml] 7.35E+08
D10 [nm] 102
D50 [nm] 141
D90 [nm] 264
Valid tracks 2800
Merged
Sample 106_treatent_1
Mode [nm] 178.5
Mean [nm] 258
Standard deviation [nm] 157
Concentration [particles/ml] 5.84E+07
D10 [nm] 118
D50 [nm] 184
D90 [nm] 407
Valid tracks 438
Concentration [particles/ml]
Size [nm]

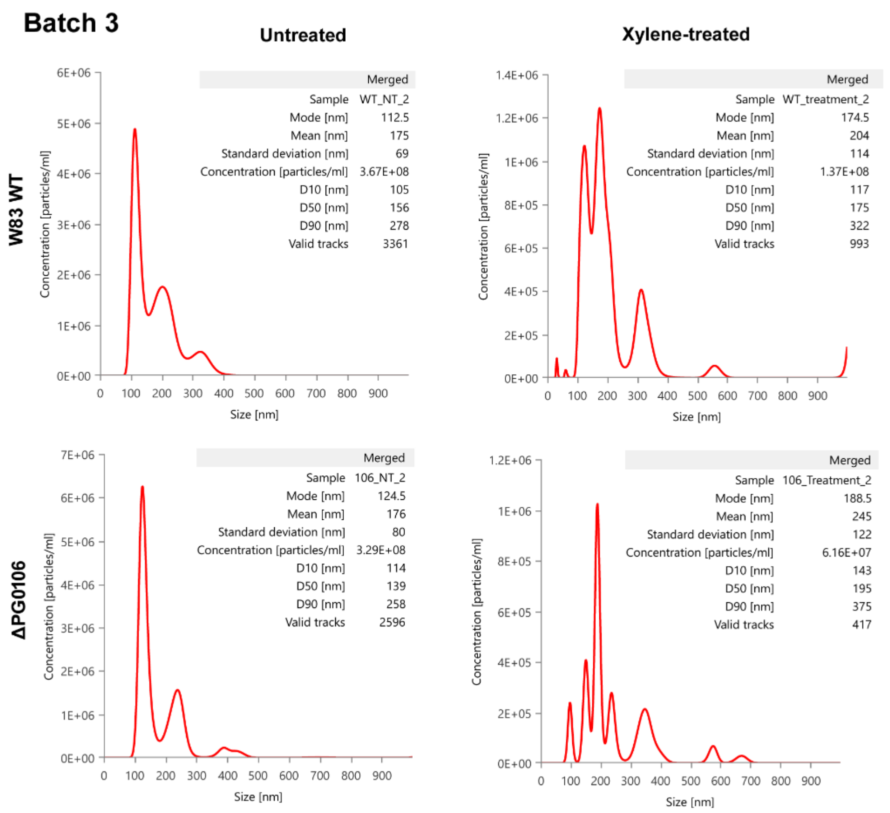


**Figure S3.** Nanoparticle Tracking Analysis (NTA) profiles of density gradient–purified OMVs from WT W83 and ΔPG0106 measured using a Nanosight Pro (Malvern Panalytical). Three independent OMV preparations (batches) were analyzed, and particle size distributions and concentrations were determined for both untreated and xylene-treated samples.

**Figure S4.**

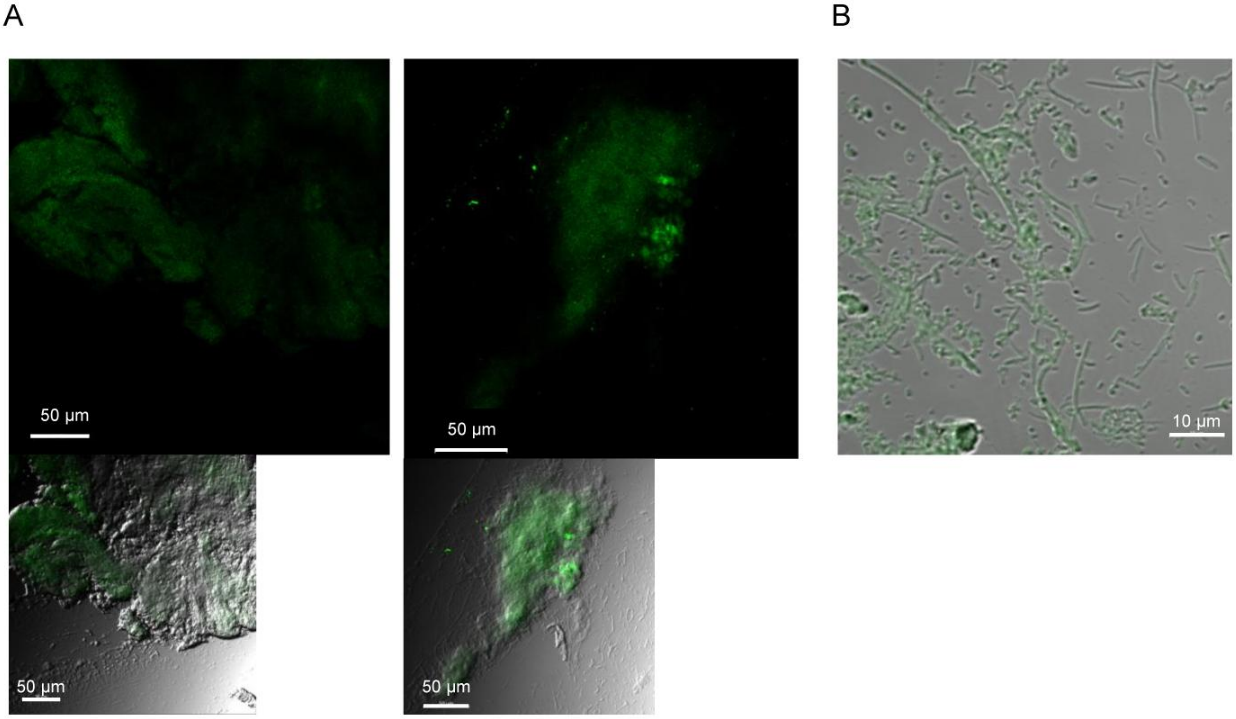


**Figure S4.** Negative control of periodontal plaque of immunofluorescence staining without primary antiserum but using IgM-AF488 only. **(A)** Zoom-out views. **(B)** A Zoom-in view.

**Figure S5.**

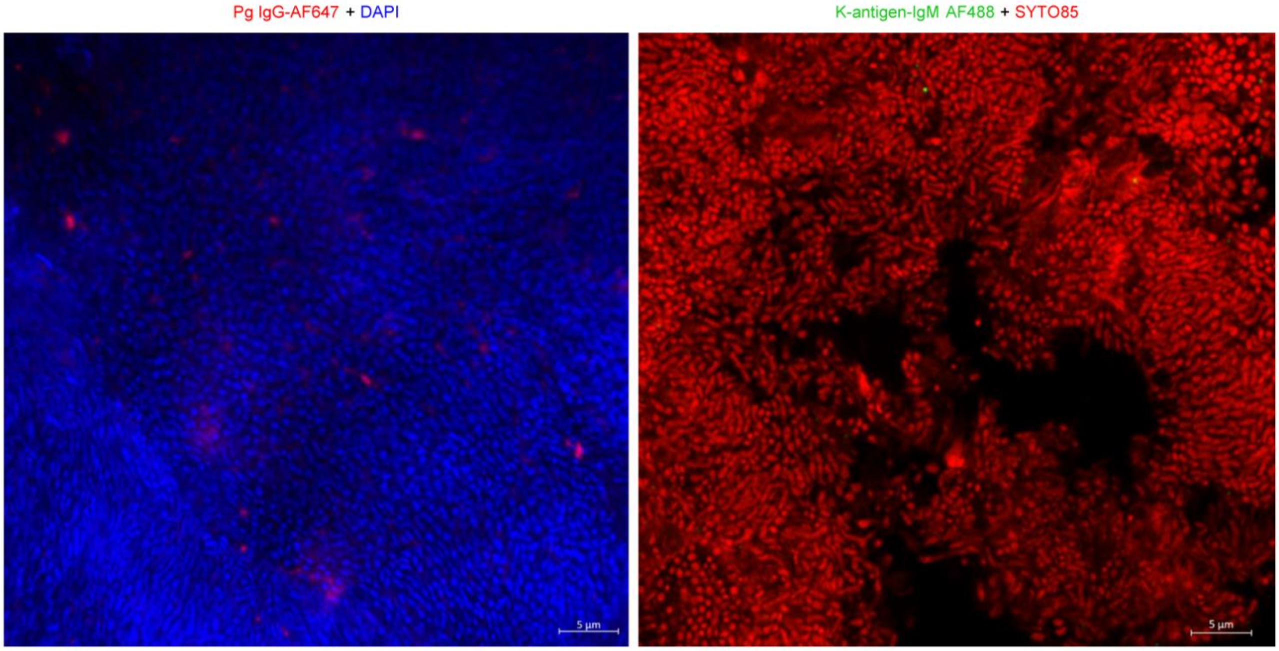


**Figure S5.** Plaque samples obtained from healthy individuals incubated with Pg IgG-AF647 (red) and stained with DAPI (blue)(left); and with the pre-adsorbed antibodies and anti-IgM-AF488 (green) for K-antigen detection and stained with SYTO85 (red) (right).

### Supplementary Methods

#### Bacterial strains and culture conditions

For subsurface grown cells, *P. gingivalis* cells from a BAPHK plate were inoculated in TSBHK medium and grown for 24 hrs. The culture was then sub-cultured at a 1:10 dilution into fresh TSBHK and grown overnight to an $OD_{600}$ of 1.0 and 100x concentrated by centrifugation. A microliter of culture was injected onto the plastic surface below the soft agar. Subsurface grown cells were observed 72 hrs after anaerobic incubation.

#### Immunofluorescent staining and confocal microscopy

*P. gingivalis* cells or plaque samples were fixed with 2–4% PFA and dried on glass slides for at least 1 hr. The dried samples were then covered with blocking solution consisting of PBS, 0.1% Tween-20, and 1× Pierce Clear Milk Blocking Buffer (Thermo Fisher Scientific) for 30 mins. The sample was incubated with blocking solution with the corresponding antiserum for 30 mins. For *P. gingivalis* detection, Alexa Fluor 647 labeled (Invitrogen) W83 antiserum (1:200) was used without the need of secondary antibodies. For K-antigen (pAds106 antiserum), 1:200 of pre-adsorbed antiserum was used for incubation for 30 min and then washed 3 times with PBS-T (PBS + 0.1% Tween-20). And then, the samples were incubated with secondary anti-rabbit IgM-AF488 antibody (Abcam) (1:500) in blocking buffer for 30mins. The samples were washed 3 times in PBS-T and covered by a cover slip for laser confocal microscopy analysis. SYTO9, SYTO85 or DAPI (Invitrogen) was added during secondary antibody incubation if necessary. Labeled sample was observed with Zeiss LSM 880 confocal upright microscope equipped with a 32-channel GaAsP Airyscan detector for super-resolution capabilities. Transmitted light images were captured using the T-PMT detector. Data shown are representative of results obtained from at least two independent experiments.

#### Outer membrane vesicle (OMV) purification

Bacterial strains in this study were cultured at 37°C in an anaerobic chamber for 16 hrs in TSBHK. After culturing, the optical density ($OD_{600}$) was adjusted to 1.0–1.2. OMVs were purified as described previously with minor modifications (45, 46). Bacterial culture supernatants were collected by centrifugation (10,000 ×*g*, 4°C, 30 mins) using a F12-6x500 rotor (Thermo Fisher Scientific) and concentrated (4,000 ×*g*, 4°C, 30 mins) using a 100 kDa cut-off centrifugal filter (Merck Millipore). The filtrates were washed with PBS and subjected to ultracentrifugation (120,000 ×*g*, 4°C, 2 hrs) using a Type 45 Ti rotor (Beckman Coulter). The crude OMV pellet was resuspended in 40% OptiPrep (0.9 mL; Sigma), overlaid with 35% OptiPrep (1.55 mL) and 10% OptiPrep (1.55 mL), and subjected to buoyant density gradient ultracentrifugation (160,000 × *g*, 4°C, 4 hrs) using an SW 60 Ti rotor (Beckman Coulter). Density fractions (400 µL each) were obtained from the top of the gradient (#1–10), and fraction #5 was washed with PBS (3.6 mL) by ultracentrifugation (120,000 ×*g*, 4°C, 2 hrs) using an SW 60 Ti rotor (Beckman Coulter). The purified OMVs were dissolved in 200 µL of PBS and stored at -80°C until use. Protein concentration of the OMVs was quantified using the bicinchoninic acid (BCA) assay (Thermo Fisher Scientific).

**Immunoblot analysis**

For K-antigen detection, cells ($OD_{600}$ = 1, in PBS) or OMVs corresponding to 10 µg of total protein were resuspended in an equal volume of 4X Laemmli sample buffer (Bio-Rad) and boiled for 5 min. For immunoblot analysis, lysates were separated and transferred onto a PVDF membrane. The membrane was blocked in Phosphate-buffered containing 0.1% Tween 20 (PBS-T) with 1X Pierce Clear Milk Blocking Buffer (Thermo Fisher Scientific). Primary pAds106 antiserum was added at 1:1000 dilution and incubated with the membrane for 1 hr with rocking at room temperature. The membrane was washed and incubated with HRP-conjugated anti-rabbit IgM (Abcam) at 1:5000 dilution for 1 hr with rocking at room temperature. The membrane was washed in PBS-T before detection using SuperSignal West Pico Chemiluminescent Substrate (Thermo Fisher Scientific). In parallel, the same samples were subjected to silver staining using the Pierce Silver Stain for Mass Spectrometry Kit (Thermo Fisher Scientific). Data shown are representative of results obtained from at least two independent experiments.

**Enzyme-linked immunosorbent assay (ELISA)**

OMVs and bacterial cell lysates were coated onto High binding 96-well plate (Greiner Bio-One) at 4°C for 16 hrs. The antigen-coated plate was washed four times with TBS-T (Tris-buffered saline containing 0.1% Tween 20) and blocked with 1% BSA in PBS at room temperature for 1 hr. After four additional washes with TBS-T, the plate was incubated with anti-*P. gingivalis* serum (1:200 dilution) at room temperature for 2 hrs, followed by four washes with TBS-T. To detect antigen-binding IgG and antigen-binding IgM, HRP-conjugated goat anti-rabbit IgG (1:20,000 dilution) and HRP-conjugated Goat Anti-Rabbit IgM (Abcam; 1:5,000 dilution) were applied, respectively. After incubation at room temperature for 20 mins, the plate was washed four times with TBS-T. Signal development was performed using 1-Step™ TMB ELISA Substrate Solutions (Thermo Fisher Scientific), and the reaction was stopped with 2 N $H_2SO_4$. The absorbance was measured at 450 nm using a SpectraMax (Molecular Devices).

**BATH assay**

Purified OMVs were first quantified by nanoparticle tracking analysis (NTA) using a Nanosight Pro instrument (Malvern Panalytical), and the particle concentration was adjusted to ~$10^8$ particles/mL. OMV suspensions (500 µL) in PBS were mixed with xylene (400 µL) and vigorously vortexed for 1 min and the mixture was then centrifuged at 1,000 × *g* for 1 min to achieve phase separation. The upper organic (xylene) phase was carefully removed, and 300 µL of the lower aqueous phase was collected and diluted with an equal volume (300 µL) of PBS. Untreated OMVs diluted in parallel and xylene-treated samples with the same final dilution were analyzed by Nanosight Pro to determine the number of particles remaining in the aqueous phase. The percentage of hydrophobicity was calculated as: 100% × (1 - number of OMVs in the xylene-treated sample divided by number of OMVs in the untreated sample).

**Alcian blue staining**

Isolated OMVs were mixed with SDS loading buffer, boiled for 5 mins, separated by SDS–PAGE. The gel was incubated in Alcian blue solution (Sigma) at room temperature with gentle agitation for 1 hr. Excess dye was removed by destaining in 3% acetic acid until the background was sufficiently reduced. Stained polysaccharides were visualized as blue bands and documented by photography on a lightbox.

**Capsular polysaccharide extraction**

Capsular polysaccharides were extracted using a phenol–water–based method with a commercially available kit (Boca Scientific, Cat#: 17141) following the manufacturer's instructions. Residual proteins were removed by treatment with proteinase K (25 µg) at 50 °C for 1 hr. LPS contaminants were eliminated by incubation in 2% acetic acid at 100 °C for 1 hr, as previously described (35).
